# Stanford Accelerated Intelligent Neuromodulation Therapy for Treatment-Resistant Depression (SAINT-TRD)

**DOI:** 10.1101/581280

**Authors:** Eleanor J. Cole, Katy H. Stimpson, Brandon S. Bentzley, Merve Gulser, Kirsten Cherian, Claudia Tischler, Romina Nejad, Heather Pankow, Elizabeth Choi, Haley Aaron, Flint M. Espil, Jaspreet Pannu, Xiaoqian Xiao, Dalton Duvio, Hugh B. Solvason, Jessica Hawkins, Austin Guerra, Booil Jo, Kristin S. Raj, Charles Debattista, Jennifer Keller, Alan F. Schatzberg, Keith D. Sudheimer, Nolan R. Williams

## Abstract

**Background:** Current treatments for depression are limited by suboptimal efficacy, delayed response, and frequent side effects. Intermittent theta-burst stimulation (iTBS) is a non-invasive brain stimulation treatment that is FDA-approved for treatment-resistant depression (TRD). Recent methodological advancements suggest iTBS could be improved through 1) treating with multiple sessions per day at optimally-spaced intervals, 2) applying a higher overall pulse-dose of stimulation and 3) precision targeting of the left dorsolateral prefrontal cortex (L-DLPFC) to subgenual anterior cingulate cortex (sgACC) circuit. We examined the feasibility, tolerability, and preliminary efficacy of an accelerated, high-dose, resting-state functional connectivity MRI (fcMRI)-guided iTBS protocol for TRD termed ‘Stanford Accelerated Intelligent Neuromodulation Therapy (SAINT)’.

**Methods:** Twenty-one participants with TRD received open-label SAINT. FcMRI was used to individually target the region of L-DLPFC most anticorrelated with sgACC. Fifty iTBS sessions (1800 pulses per session, 50-minute inter-session interval) were delivered as 10 daily sessions over 5 consecutive days at 90% resting motor threshold (adjusted for cortical depth). Neuropsychological testing was conducted before and after SAINT.

**Results:** Nineteen of 21 participants (90.48%) met criteria for remission (≤10 on the Montgomery-Åsberg Depression Rating Scale) immediately after SAINT. Neuropsychological testing demonstrated no negative cognitive side-effects. There were no seizures or other severe adverse events.

**Discussion:** Our accelerated, high-dose, iTBS protocol with fcMRI-guided targeting (SAINT) was well tolerated and safe. Efficacy was strikingly high, especially for this treatment-resistant population. Double-blinded sham-controlled trials are required to confirm the high remission rate found in this initial study.

**Trial registration:** ClinicalTrials.gov NCT03240692

## Introduction

Depression is the leading cause of disability worldwide, and approximately 800,000 suicides are completed each year (1–3). Current FDA-approved antidepressant treatments do not achieve remission in the majority of patients with treatment-resistant depression (TRD) (4–6), are limited by tolerability (7) and have extended treatment durations, which do not match the imminent risk to suicidal patients (8–10). New antidepressant treatments are needed that are safe, tolerable, rapid-acting, durable, and more effective than current interventions.

Repetitive transcranial magnetic stimulation (rTMS) delivered to the left dorsolateral prefrontal cortex (L-DLPFC) is an FDA-approved non-invasive brain stimulation technique for TRD (11). rTMS involves passing an electrical current through a magnetic coil placed superficial to the scalp, producing a high-intensity magnetic field that passes through the scalp, skull and meninges to excite neuronal tissue (12). Repeated high-frequency excitation of the same brain region results in strengthening of synapses through a process known as long-term potentiation (LTP) (13, 14), causing changes in functional connectivity (13, 15). The antidepressant mechanism of rTMS is hypothesized to be mediated in part through indirect inhibitory functional connectivity from L-DLPFC to sgACC (15–17).

A more efficient form of rTMS, known as intermittent theta-burst stimulation (iTBS), has been developed, which has significantly shortened the duration of treatment sessions from 37 minutes to 3 minutes (18) and produces equivalent antidepressant responses (19, 20). FDA-approved rTMS and iTBS courses involve daily stimulation sessions (600 iTBS pulses) for 6 weeks, achieving remission in 32% of patients and response in 49% (19). Studies suggest that the efficacy of iTBS could be improved by optimized iTBS session spacing, accelerated delivery (21–24), higher overall pulse-doses (10, 25, 26) and individualized targeting (15, 27).

We implemented all of these potential improvements on rTMS treatment of depression into an accelerated, high-dose, iTBS protocol using functional connectivity magnetic resonance imaging (fcMRI)-guided targeting. This protocol includes 5 consecutive days of 10-daily iTBS sessions (1800 pulses per session) delivered to the region of the L-DLPFC that is most functionally anticorrelated with the sgACC in each individual (28). Individualized fcMRI-guided targeting, optimized intersession interval spacing, accelerated delivery and high pulse-dose were predicted to collectively result in higher response and remission rates than current FDA-approved TMS protocols. This protocol was termed ‘Stanford Accelerated Intelligent Neuromodulation Therapy (SAINT)’ to distinguish this protocol from other attempts at accelerating TMS protocols without individualized targeting, optimized intersession spacing or high pulse-dose (29, 30). Our incipient investigation of SAINT demonstrated surprisingly high efficacy in a small cohort of participants with extremely severe and treatment refractory depression (28)^a^. The study we describe herein builds upon our initial report by testing SAINT in a larger and more generalizable cohort of participants with TRD to examine the feasibility, safety and preliminary efficacy of this approach.

## Methods

### Participants

Participants were required to be currently experiencing a non-psychotic, non-atypical major depressive episode as part of either bipolar II disorder or major depressive disorder (MDD) as defined by DSM-5 criteria and not responded to at least 1 antidepressant medication. At the time of screening, participants were required to have a Hamilton Depression Rating Scale 17-item (HDRS-17) score of 20 or higher, a negative urine drug screen and negative urine pregnancy test if female. Participants were excluded if they had any contraindications to rTMS such as history of seizures, metallic implants in the head, cardiac pacemakers or a neurological disorder. Participants were recruited through the Depression Research Clinic at Stanford University, study advertising and clinic referrals.

Twenty-three participants (aged 19-78, 13 female) were recruited for this study. One participant was screened out post-enrollment for having a very high motor threshold (>90% machine output) and 1 participant with a history of multiple prior therapeutic intolerances (anxiety leading to early discontinuation of intravenous ketamine infusions and conventional rTMS) dropped out after the first day of stimulation due to anxiety. This resulted in a final sample of 21 participants (aged 19-78, 12 female). Nineteen participants had a diagnosis of MDD, and 2 participants had a diagnosis of bipolar II disorder currently in a depressive episode (>1 year). See Table 1 for demographic information and treatment history. Participants were required to maintain their antidepressant regimen throughout study enrollment (see Supplementary Table 1 for medications taken during enrollment).

**Table 1:**
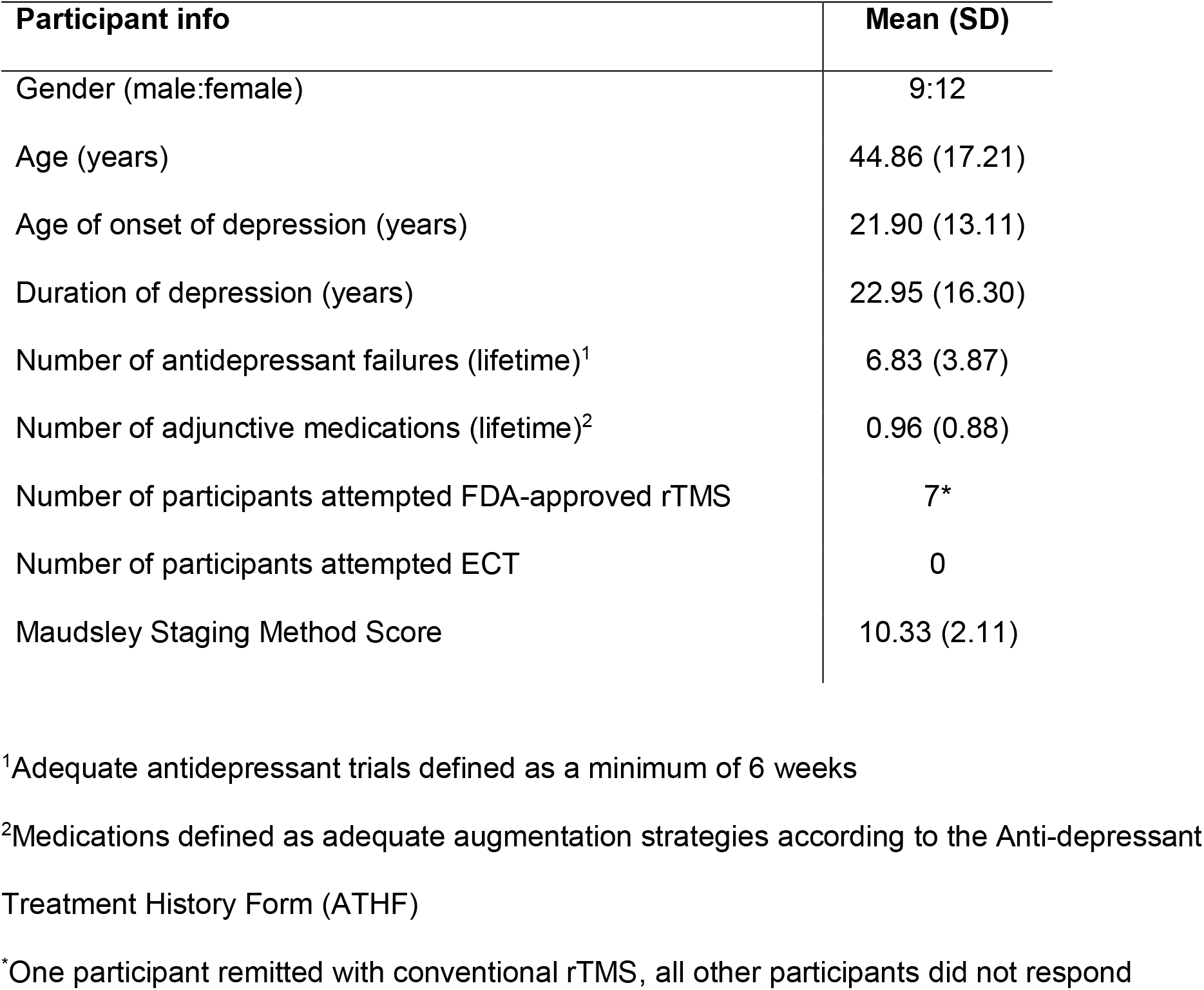
Demographic information and treatment history for all participants (n=21)

| **Participant info** | **Mean (SD)** |
| --- | --- |
| Gender (male:female) | 9:12 |
| Age (years) | 44.86 (17.21) |
| Age of onset of depression (years) | 21.90 (13.11) |
| Duration of depression (years) | 22.95 (16.30) |
| Number of antidepressant failures (lifetime)<sup>1</sup> | 6.83 (3.87) |
| Number of adjunctive medications (lifetime)<sup>2</sup> | 0.96 (0.88) |
| Number of participants attempted FDA-approved rTMS | 7* |
| Number of participants attempted ECT | 0 |
| Maudsley Staging Method Score | 10.33 (2.11) |

<sup>1</sup>Adequate antidepressant trials defined as a minimum of 6 weeks
<sup>2</sup>Medications defined as adequate augmentation strategies according to the Anti-depressant Treatment History Form (ATHF)
\*One participant remitted with conventional rTMS, all other participants did not respond

### Functional magnetic resonance imaging (fMRI)

Before the stimulation course, each participant had both structural MRI and resting-state fMRI scans. All MRI scans were acquired using a 3-tesla GE Discovery MR750 scanner with a 32- channel imaging coil at the Center for Cognitive and Neurobiological Imaging at Stanford, using a 3x accelerated multiband imaging sequence with a repetition time of 2 seconds. During the 8-minute resting state scans, participants were instructed to let their minds wander, avoid repetitive thoughts, keep their eyes open and their attention focused on a central fixation point.

### Stanford Accelerated Intelligent Neuromodulation Therapy

A MagVenture MagPro X100 (MagVenture A/S, Denmark) system was used to deliver sessions of iTBS; 60 cycles of 10 bursts of 3 pulses at 50Hz were delivered in 2-second trains (5Hz) with an 8-second inter-train interval. Stimulation sessions were delivered hourly (21–23). Ten sessions were applied per day (18,000 pulses/day) for 5 consecutive days (90,000 pulses in total). Stimulation was delivered at 90% resting motor threshold (rMT; 30–32). A depth correction was applied (34) to consistently achieve 90% rMT at the depth of the functional target. Stimulation was never delivered above 120% rMT for safety (35). The Localite Neuronavigation System (Localite GmbH, Sankt Augustin, Germany) was used to position the TMS coil over the individualized stimulation target. See Figure 1 for differences between SAINT and the FDA-approved iTBS protocol.

**Figure 1:**
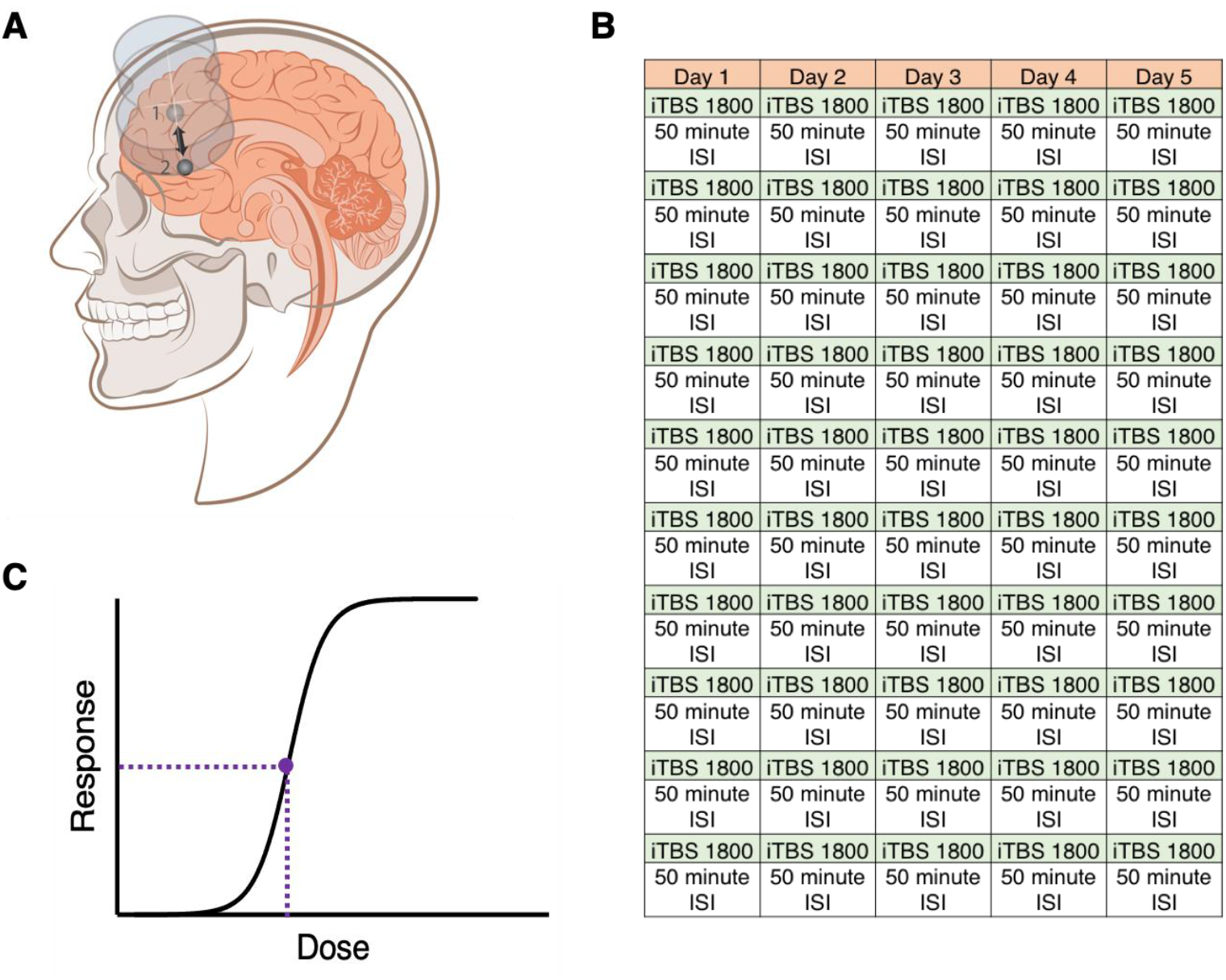
This figure illustrates the 3 factors that make SAINT different from current FDA-approved iTBS protocols: Individualized targeting (A), accelerated delivery of hourly sessions (B) and high pulse-dose [both per session (B) and overall (C)]. The FDA-approved iTBS protocol involves 1 daily stimulation session of 600 pulses to the L-DLPFC at an intensity of 120% RMT for 6 weeks. The location of the L-DLPFC is located using scalp measurements. In our study, A) we used fMRI scans to target the region of the L-DLPFC (1) with functional connectivity that was most anticorrelated with the sgACC (2). This was done, because previous neuroimaging findings suggest the higher the anticorrelation between the stimulated region of the L-DLPFC and the sgACC, the better the clinical outcome (15, 27, 92). B) 10 sessions of 1800 pulses per day. 1800 pulses was chosen as this is the only pulse-dose that has been explored in a blinded iTBS trial (60). Additionally, 1800 pulses has been shown to produce long-lasting changes in cortical excitability (93) and optimally produce intended cellular changes (94). C) The highest overall pulse-dose of any published study to date. A dose-response curve has not been created for TMS. A previous study showed 61% of non-responders to rTMS responded with further treatment (26) suggesting that FDA-approved protocols are under-dosing. Our SAINT protocol delivered five-times the FDA-approved pulse-dose. See supplementary material for more information regarding protocol development.

### Clinical assessments

Before and after SAINT, depressive symptoms and suicidal ideation were assessed using clinical and self-report assessments [HDRS-17, Montgomery-Åsberg Depression Rating Scale (MADRS), Columbia-Suicide Severity Rating Scale (C-SSRS, suicidal ideation subscale), Beck Depression Inventory-II (BDI-II)]. At the end of each day of stimulation (10 sessions), depressive symptoms were assessed using the Hamilton Depression Rating Scale 6-item (HDRS-6). The Young Mania Rating Scale (YMRS) was completed daily to assess for hypomania (36).

A neuropsychological test battery was administered before and after SAINT to capture any neurocognitive side effects. The Hopkins Verbal Learning Test – Revised (HVLT-R) (37), the Brief Visuospatial Memory Test – Revised (BVMT-R) (38), subtests from the Wechsler Adult Intelligence Scale (4th Ed.; WAIS-IV) (39) and several tests from the Delis Kaplan Executive Function System (D-KEFS) (40) were used. See supplementary material for detailed information about the neuropsychological test battery.

### fMRI analysis for target generation

Personalized L-DLPFC targets were generated for each participant using the baseline resting-state scan. All analyses were conducted in a participant’s own brain space. Resting-state scans were pre-processed according to typical methods using Statistical Parametric Mapping (SPM12) software. The resting-state scans were motion-corrected and resliced. T1-weighted structural scans were then co-registered with the resting-state scans. Next, the estimation parameters to warp the T1-weighted structural image into Montreal Neurological Institute (MNI) space were calculated using SPM segmentations based on tissue probability maps. These normalization parameters were inverted and applied to MNI space regions of interests (ROIs) for the L-DLPFC (Brodmann area 46) and the sgACC (BA25) to map these ROIs onto the individual participant’s brain. The participant-space ROIs were then resliced, smoothed, and binarized to match the dimensions of the resting-state scans.

The participant-space ROI for the L-DLPFC formed the search area for the optimal TMS coil placement. Two separate algorithms were used to determine coil placement. The first algorithm sorted each of the L-DLPFC and bilateral sgACC voxels into functional sub-units using a hierarchical agglomerative clustering algorithm. Median time series were then created for each functional subunit, and the correlation coefficients were calculated between all median time series extracted from all functional subunits of the L-DLPFC and sgACC. The second algorithm determined the optimal L-DLPFC subunit to target based on 3 factors: the net correlation/anticorrelation of the L-DLPFC subunit with sgACC subunits, the size of the subunit and the spatial concentration of the subunit. See supplementary methods for more details on these algorithms. Three-dimensional maps of the whole brain correlation coefficient of the selected L-DLPFC subunit were then created and used to target the coil placement using the Localite TMS Navigation software (Localite GmbH, Sankt Augustin, Germany). See Figure 2 for the individual target locations.

**Figure 2:**
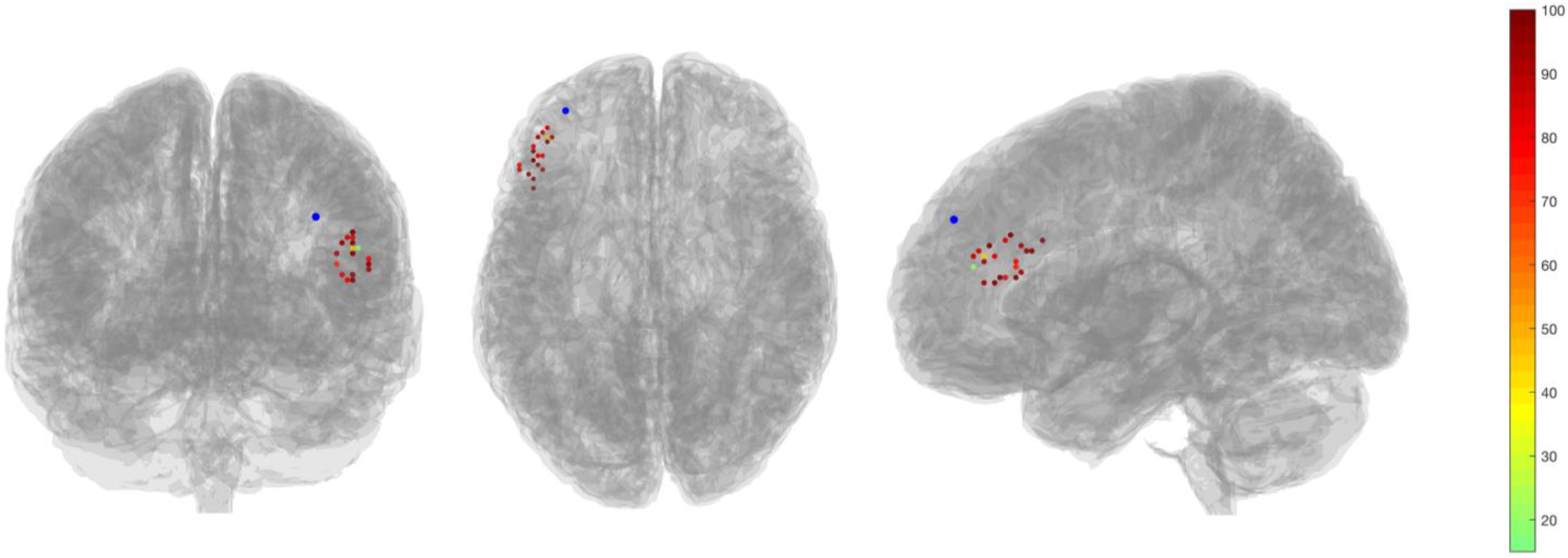
Individual target locations used in this study in comparison to an average F3 coordinate (−35.5, 49.4, 32.4) shown in blue (95). The colors of the targets represent the % change in MADRS score with dark red indicating greater change in MADRS score. Mean distance from F3 was 25.18mm (SD=6.15).

### Clinical outcome analysis

All statistical analyses were conducted using IBM SPSS Version 22. The level of statistical significance was set at *p*=0.05. Missing data were not imputed. Statistical analyses were planned independently by BSB and BJ and reviewed by AFS and NRW.

Our primary outcome measure was change in MADRS scores from baseline to immediately following SAINT, and MADRS scores were used to calculate response and remission rates. Reductions in HDRS-17, HDRS-6 and BDI-II scores were used as secondary outcome measures of depression severity. Response was defined as ≥50% reduction in these scales. Remission was defined as MADRS score ≤10 (41), ≤7 on HDRS-17 (42), ≤4 on the HDRS-6 (43) and ≤12 on the BDI-II (44). A floor effect of SAINT treatment was observed across all scales, and initial linear mixed models produced residuals that were not normally distributed (Shapiro-Wilk). Thus, changes in MADRS, HDRS-17, and BDI-II were assessed with generalized linear mixed models (GLMM) that used a compound symmetry covariance structure, Satterthwaite approximation of degrees of freedom, and robust estimation of coefficients to handle violations of model assumptions. Fixed effects of time and a treatment history of non-response to conventional rTMS and their interaction were assessed. All post-hoc pairwise comparisons were Bonferroni-corrected.

Daily HDRS-6 scores were used to calculate the number of days of stimulation required to reach responder criterion (≥50% reduction from baseline on HDRS-6) and remission criterion (≤4 on HDRS-6). Kaplan-Meier survival analysis using the Breslow test of equality of survival distributions was used to determine if there were significant differences in the number of days to reach response and remission criteria for participants who had a treatment history of conventional rTMS non-response compared to those who did not.

Suicidality was assessed using the suicidal ideation subscale of the C-SSRS, item 3 of the HDRS-17, and item 10 of the MADRS. Response was defined as ≥50% reduction in these scores from baseline, and remission was defined as a score of 0. Response was only calculated if the baseline score was >0. Scores were ordinal and changes in scores were assessed with generalized linear models (GLM) with a multinomial link, compound symmetry covariance structure, Satterthwaite approximation of degrees of freedom, and robust estimation of coefficients to handle violations of model assumptions.

Scores on the neuropsychological tests before and after SAINT were compared using paired t-tests. The data for the HVLT-R total score, the BVMT-R-Delayed Recall and the number of rule violations on the D-KEFS Tower Test violated the assumption of normality, so non-parametric (Wilcoxon signed-rank) tests were used to evaluate SAINT-induced changes in performance on these 3 measures. Neuropsychological test data were available for 17 participants.

### Research ethics and approval

All research procedures were conducted in accordance with the ethical standards outlined in the Declaration of Helsinki. This study was approved by the Stanford University Institutional Review Board. All participants provided written informed consent before taking part in any study procedures.

## Results

### Safety

No serious adverse events occurred. As stated in the Methods, 1 participant with a history of multiple prior therapeutic intolerances (anxiety leading to early discontinuation of intravenous ketamine infusions and conventional rTMS) dropped out after the first day of stimulation due to anxiety. The only side-effects other participants reported were fatigue and some discomfort at both the stimulation site and in the facial muscles during stimulation. The neuropsychological test battery showed no negative cognitive side-effects following SAINT. Performance significantly improved on measures of cognitive inhibition [D-KEFS color-word inhibition task; t(16)=4.92, *p*<0.000, d=1.19; D-KEFS color-word inhibition switching task; t(16)=3.77, *p*=0.002, d=0.91]. These improvements survived correction for multiple comparisons (Bonferroni-corrected significance level *p*<0.004). There were no significant changes on any of the other neurocognitive tasks; see supplementary material.

### Depression symptoms

GLMM analysis revealed a significant effect of time (*F*_3,9_=90.423, *p*<0.001) on mean MADRS scores with all follow-up time points being significantly lower than baseline (Bonferroni-corrected pairwise comparisons, *p*<0.01). These results were recapitulated for the HDRS-17 (*F*_3,12_=51.765, *p*<0.001) and the BDI-II (*F*_3,19_=19.040, *p*<0.001). The response rate (≥50% reduction in MADRS) was 90.48%, and all responders were in remission (MADRS score ≤10). Results were similar across all clinical assessments (see Table 2). One month following SAINT, 70% of participants continued to meet responder criteria (see Supplementary Table 2 for response and remission rates at 1 month).

**Table 2:**
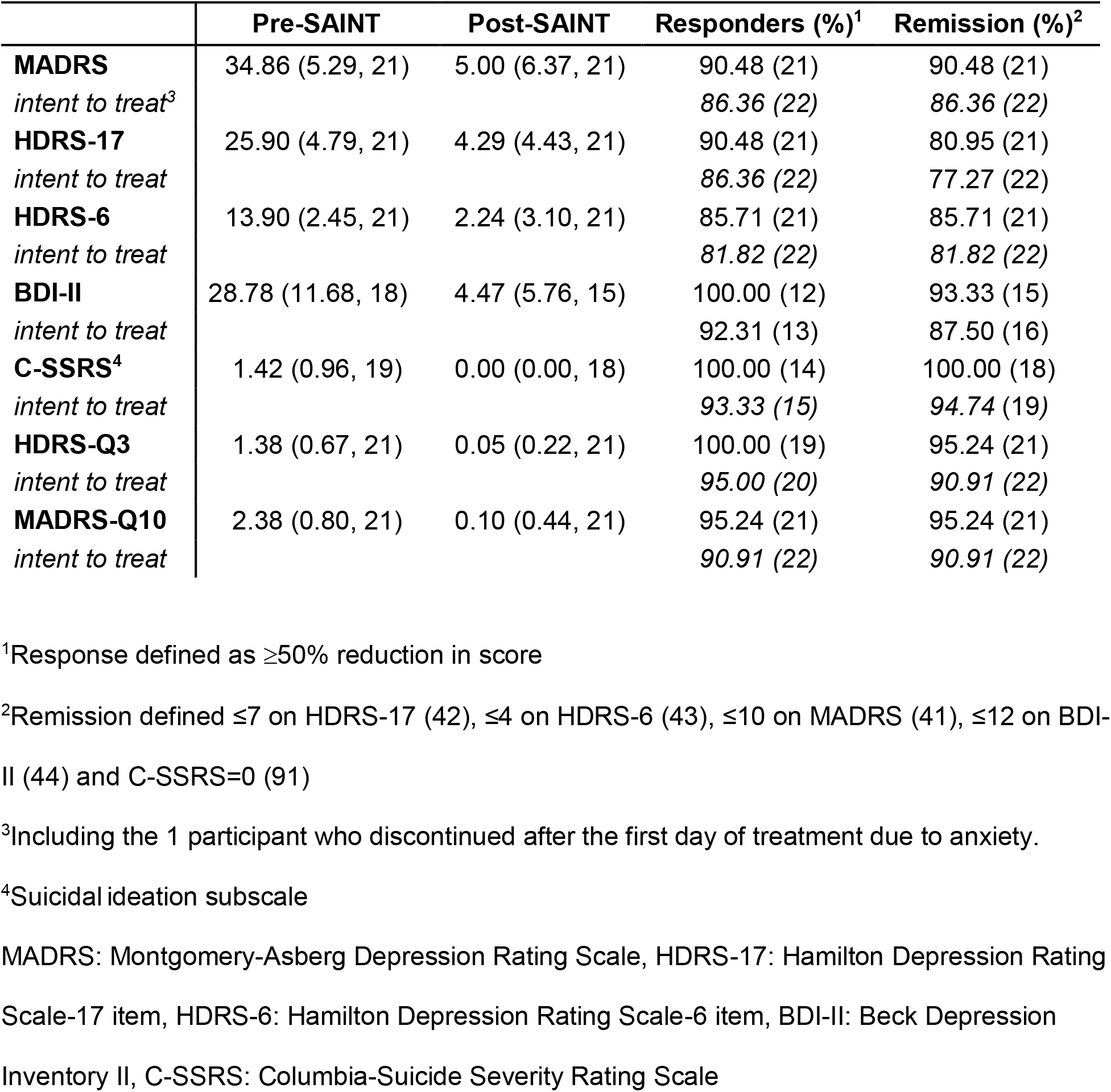
Clinical assessment scores for all participants immediately following SAINT; mean (SD, n) or % (n)

A hypothesis of SAINT is that conventional rTMS delivers insufficient cumulative stimulation to induce response and remission from depression for some patients. We tested this in part by including participants who had a history of non-response to conventional rTMS (rTMS non-responders, n=6). When comparing the non-responder group to the rest of the participants, we found that conventional rTMS non-responders had similar MADRS scores at baseline. TMS non-responders had greater mean MADRS scores at every time point after SAINT; however, neither the main effect of group (GLMM, *F*_1,4_=5.695, *p*=0.072, n.s.) nor the group x time interaction reached statistical significance (*F*_3,9_=1.077, *p*=0.405, see Supplementary Table 3), indicating that participants with a history of conventional rTMS non-response had a similar treatment effect as the other participants.

### Days of treatment until response and remission

The mean number of days of SAINT completed until participants met the response criterion (≥50% reduction in HDRS-6 score) was 2.30 (SD=1.13, ~23 10-minute treatments, n=20, daily HDRS-6 scores missing for 1 participant), and the mean number of days to achieve remission (HDRS-6 score ≤4) was 2.63 (SD=1.21, ~26 10-minute treatments, n=19, 1 participant did not achieve remission by the HDRS-6 criterion). See Figure 3 for percentage change in HDRS-6 score with each day of stimulation.

Kaplan-Meier survival analysis revealed that participants who had previously not responded to a 6-week rTMS treatment course (n=6) required more days of treatment to achieve responder criterion [***X***^2^=4.359, *p*=0.037, mean 3.00 days (SD=0.63), ~30 10-minute treatments] and this approached statistical significance for remission criterion as well [***X***^2^=3.558, *p*=0.057, mean 3.20 days (SD=0.84), ~32 10-minute treatments, n=5, 1 participant did not meet criteria for remission). See Figure 3.

**Figure 3:**
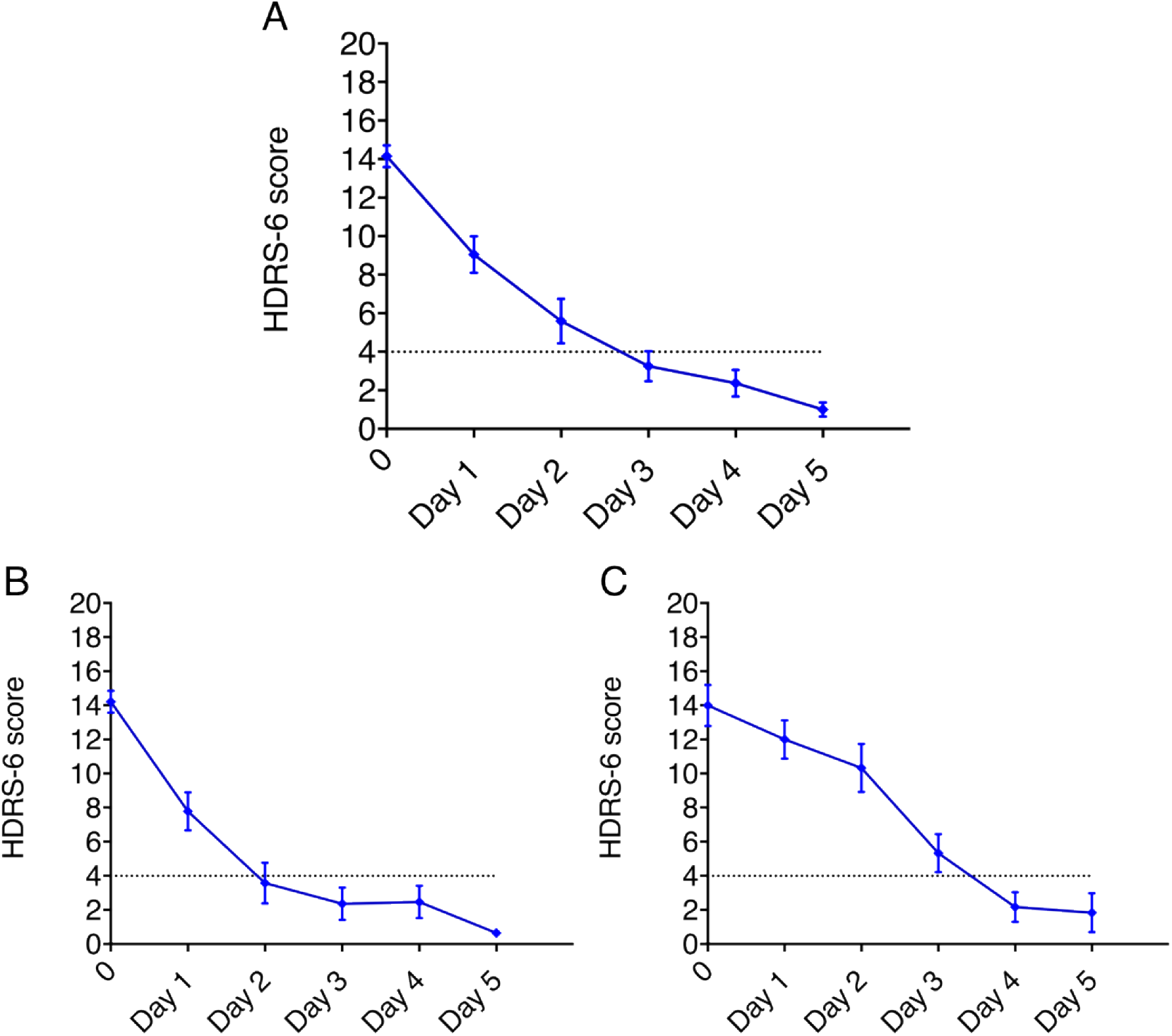
Mean Hamilton Depression Rating Scale score (6-item version, HDRS-6) with each day of stimulation for A) all participants B) participants excluding TMS non-responders, C) TMS non-responders. Dotted lines indicate remission criteria and error bars represent standard error.

### Suicidality scales

Of the 21 total participants at the time of screening, 19 reported some degree of suicidality on the C-SSRS, 20 reported suicidality on item 3 of the HDRS-17 and all 21 reported suicidality on item 10 of the MADRS. Changes in suicidality scales were assessed with GLM with a multinomial link. Following SAINT there were significant reductions in the C-SSRS (GLM, X^2^(1)=16.398, *p*<0.001), item 3 of the HDRS-17 (GLM, X^2^(3)= 31.057, *p*<0.001), and item 10 of the MADRS (GLM, X^2^(3)=46.855, *p*<0.001) at all follow up time points (X^2^, *p*<0.001). See Table 2. One month following SAINT, 80-100% of participants continued to be remitted on these measures (see Supplementary Table 2).

## Discussion

The aim of this study was to examine the safety, feasibility and preliminary efficacy of an accelerated, high-dose iTBS, fcMRI-guided treatment protocol (SAINT) for TRD. We found that SAINT significantly reduced depressive symptoms and suicidal ideation in patients with severe TRD within 5 days, without negative cognitive side-effects or other adverse events. The remission rate we observed is substantially higher than remission rates for open-label studies using standard FDA-approved rTMS treatment protocols (19, 45, 46). This is also higher than the remission rates that have been reported for electroconvulsive therapy (ECT) treatment (~48%) (5) and for ketamine treatment of TRD (31%) (47). See Supplementary Table 4 for remission rates for interventions for TRD. This remission rate was observed despite the inclusion of participants who had previously not responded to FDA-approved rTMS. The high efficacy of SAINT may be due to optimally-spaced iTBS sessions, accelerated delivery, high pulse-dose, individualized targeting or a combination thereof.

The high efficacy of our SAINT protocol complements evidence from basic neuroscience research and human physiology data, which suggest that multiple optimally-spaced daily iTBS sessions have an enhanced effect compared to the same number of single daily sessions (21–23, 48–51). Multiple optimally-spaced stimulation sessions have also been shown to produce accumulating non-linear improvements in clinical symptoms (52, 53). The duration of inter-session intervals appears to be vitally important as stimulation sessions with inter-session intervals of 50-90 minutes have been shown to have a cumulative effect on synaptic strengthening; whereas, sessions with inter-session intervals of 40 minutes or less do not show this cumulative effect (21–23). Similarly, some studies have shown that 2 theta-burst stimulation sessions delivered to the motor cortex 15 minutes apart do not increase cortical excitability compared to a single session (51, 54). Finally, L-DLPFC activity has been shown to be correlated with sgACC activation 10 minutes after rTMS; this correlation was reduced 27 minutes following rTMS; and the correlation between these regions reached its nadir 45 minutes after rTMS (17), i.e. the hypothetically optimal time for stimulation. Taken together, these data could explain the limited efficacy of a previously reported accelerated stimulation protocol for depression, which used an inter-session interval of only 15 minutes (29, 30).

The individualized targeting method used in our study may have also contributed to the high remission rate. The L-DLPFC is a large brain area that consists of several subregions, some of which are correlated and some anticorrelated with sgACC activity (55). A retrospective study found that the degree of anticorrelation between the stimulated subregion of the L-DLPFC and the sgACC accounted for over 70% of the variance in antidepressant efficacy of rTMS (15). A recent interleaved TMS-fMRI study showed that when using individualized functional connectivity-guided targeting, stimulation propagated from the L-DLPFC to the sgACC in all participants (56). In comparison, a separate study defined the L-DLPFC anatomically (border of BA9/BA46) and stimulation propagated to the sgACC in only 44% of participants (57). A trial in healthy individuals showed that stimulating the L-DLPFC using personalized functional connectivity-guided targeting induced the desired change in functional connectivity between the L-DLPFC and the sgACC (17). Defining L-DLPFC using common techniques such as scalp-based measurements or structural MRI scans could result in stimulating a subregion of the L-DLPFC that is correlated rather than anticorrelated with the sgACC (58, 59). The limited efficacy of prior iTBS treatment protocols (30, 60) could be explained in part by the use of this anatomical target (border of BA9/BA46). By stimulating the subregion of the L-DLPFC that is most anticorrelated with the sgACC in each individual, we may have reduced this variability in signal propagation and maximized treatment efficacy.

The high efficacy of our SAINT protocol suggests that FDA-approved protocols could be under-dosing. Our protocol administered five-times the pulse-dose of the FDA-approved iTBS protocol (90,000 pulses in comparison to the standard 18,000 iTBS pulses (19)). Prior studies found that 61% of individuals who do not respond to an rTMS treatment course responded with additional rTMS treatment sessions (26), and higher pulse-doses are associated with higher efficacy (61, 62). A recent report demonstrated non-asymptotic negative linear relationships between the number of rTMS treatments and depression symptom scores (63). Finally, the possible need for a higher pulse-dose is consistent with deep brain stimulation in other neuropsychiatric disorders, where ~500,000 pulses of stimulation are delivered each day (64). This suggests that additional rTMS treatments might further reduce depression symptoms. Our SAINT protocol applies the equivalent amount of stimulation as a 6-week standard iTBS treatment protocol (18,000 pulses) each day of stimulation (19). Thirty percent of participants in our study met responder criteria after the first day of stimulation (n=6/20, daily HDRS-6 missing for 1 participant), which is equivalent to response rates for iTBS/rTMS for this treatment-resistance level (65–68). None of the prior rTMS non-responders in our study responded after a single day of SAINT (see Figure 3); whereas, 83% of these prior rTMS non-responders did respond at the end of the 5-day protocol. The need for higher stimulation doses may have contributed to the limited efficacy of previous attempts at accelerated stimulation approaches, which delivered much lower daily and total pulse-doses (29). Our study administered the highest number of TMS pulses per day and highest overall TMS pulse-dose of any study we are aware of (28).

Prior rTMS non-responders in our study required more stimulation sessions to induce a clinically significant response. It is possible that depressed individuals with a higher degree of treatment-resistance display neuroplasticity impairments (69). Thus, highly treatment-resistant individuals may require a higher pulse-dose to induce an antidepressant response, and individuals with the highest degree of treatment resistance may require maintenance iTBS therapy (70) or even an implanted cortical stimulator (71, 72) to induce and sustain antidepressant responses (26).

The combined short duration of our SAINT protocol and the apparent lack of cognitive side-effects provide potential benefits over currently available treatments. Ketamine/esketamine and ECT are currently available alternatives for rapid antidepressant response but have several limitations. Remission rates for ketamine/esketamine are substantially lower than the remission rate we observed for SAINT (73–76), approximately 11% of patients report the dissociative symptoms from ketamine as very disturbing (76), and the opioid mechanism of action may pose a potential risk for some (77). A large trial investigating the efficacy of esketamine showed a maximal 4-point average difference on the MADRS in comparison to placebo (78). Although ECT is safe and effective, its use has been limited to less than 2% of eligible patients due to restricted availability coupled with concerns regarding cognitive side-effects and stigma (79, 80). Further, ECT often takes two weeks or longer to produce remission from suicidal ideation (81).

Our study has several limitations, including an open-label design and small sample size. Without a sham-control group we cannot rule out that our results are primarily due to sham effect. However, a previous study found individuals with high treatment-refractoriness, like many of the participants in this study, show no sham response to iTBS sessions of 1800 pulses (60). Further, this study involved 500 minutes of stimulation; whereas, conventional rTMS involves 1200 minutes of stimulation time (10, 68). Thus, it would be expected that with less time in treatment there would be less of an opportunity for a sham effect than in conventional rTMS treatment. The remission rate in this study is also substantially higher than previous open-label interventions for TRD; see Supplementary Table 4. Greater sham remission rates would be expected for DBS as sham response magnitude is related to the degree of invasiveness of the procedure (82). The pooled open-label remission rate for DBS is 30% (83). The remission rate we found (90.48%) is substantially higher than this, suggesting that SAINT is either a more effective treatment or the SAINT procedure introduced an extremely high sham effect. An observational study monitoring 124 individuals with TRD receiving treatment as usual (medications, psychotherapy and ECT)showed a 3.6% remission rate after 1 year (84) demonstrating the low incidence of spontaneous remission in this population. Regardless, a sham-controlled double-blinded study is required to define the effectiveness of our protocol in comparison to an identical schedule of sham stimulation sessions.

Further methodological uncertainties include stimulation of a single brain region (85), fixed stimulation frequencies (63, 86), fixed inter-session intervals (86, 87) and the lack of state-dependent stimulation (88). Individualized stimulation frequencies may result in quicker and more durable responses (86, 89), and different cortical excitability profiles may require different inter-session intervals (87, 90). Finally, recent studies have shown that applying stimulation in particular brain states using real-time electroencephalography-triggered TMS (EEG-TMS) can increase cortical responses to stimulation (88).

In conclusion, SAINT; our high-dose, accelerated and fcMRI-guided iTBS protocol is preliminarily safe, well-tolerated, feasible, and associated with a high rate of remission from depression, despite the inclusion of participants who had previously not responded to rTMS. Our data suggest that current FDA-approved TMS protocols may be under-dosing and could potentially benefit from individualized targeting methods and accelerated delivery of a high pulse-dose via optimally-spaced sessions. The efficacy of SAINT in treating suicidal ideation and the short duration of the protocol suggest SAINT could provide a means of rapidly ensuring the safety of suicidal patients. Larger, double-blinded, sham-controlled trials are required to confirm the promisingly high response and remission rates found in this initial study.

## Author contributions

EJC ran the experiment, collected the data, analyzed the data, created the figures and tables and wrote the manuscript.

KHS set-up the experiment, ran the experiment, collected the data and edited the manuscript.

BSB designed the experiment, analyzed the data, wrote the results section and edited the manuscript.

MG ran the experiment and collected the data.

KC assisted with the design of the neuropsychological battery, performed neuropsychological assessments, collected the resulting data, and screened potential participants.

RN ran the experiment and assisted with regulatory aspects.

HP assisted with participant recruitment, participant screening and data storage.

EC conducted neuropsychological assessments.

HA conducted neuropsychological assessments.

FME supervised psychology students performing clinical assessments for this study.

JP assistant in project construction and data collection.

XX assisted with MRI data quality and analysis.

DD conducted MRI scans.

HBS provided MD coverage for the TMS sessions and subject recruitment.

JH assisted with clinical trial design and regulatory aspects.

AG created Figure 1A.

KSR provided MD coverage for the TMS sessions and subject recruitment.

CD provided MD coverage for the TMS sessions and subject recruitment.

JK assisted with the design of the neuropsychological test battery and supervised students conducting the assessments.

AFS helped with trial design, subject recruitment, and acted as unconflicted MD investigator who discussed elements of data analysis with EJC.

KDS designed and executed the algorithm used to define the optimal stimulation targets and assisted with the design of the trial.

NRW invented the stimulation methodology, designed the trial, acted as study MD, mentored the data analysis and the writing of the manuscript by EJC.

## Acknowledgments

We would like to thank all of the participants for their contributions.

This work was supported by Charles R. Schwab, the Gordie Brookstone Fund, The Marshall and Dee Ann Payne Fund, the Lehman Family, Neuromodulation Research Fund, Still Charitable Fund, Avy L. and Robert L. Miller Foundation, Stanford Psychiatry Chairman’s Small Grant, Stanford CNI Innovation Award, NIH T32 035165, NIH UL1 TR001085, Stanford Medical Scholars Research Scholarship, NARSAD Young Investigator Award and the Department of Psychiatry and Behavioral Sciences at Stanford University.

## Footnotes

a These participants are not included in this manuscript.

