## Supplementary material for "Stanford Accelerated Intelligent Neuromodulation Therapy for Treatment-Resistant Depression (SAINT-TRD)"

Neuropsychological test battery results

There were improvements on cognitive switching tasks [Delis-Kaplan Executive Function System (D-KEFS) trail making test condition 4, Alternating Letter-Number Switching; $t(15)=3.00$, $p<0.01$, $d=0.75$], problem solving tasks [D-KEFS Tower Test Total Achievement Score; $t(16)=-2.64$, $p=0.02$, $d=0.64$]; and rule violations on the D-KEFS Tower Test; $z=-2.68$, $p=0.007$, $r=0.65$]. However, due to the large number of comparisons, these improvements did not survive correction for multiple comparisons.

There were no significant differences in performance on the remaining neuropsychological tasks [Hopkins Verbal Learning Test – Revised (HVLT-R) delayed recall: $t(16)=-0.80$, $p=0.43$, $d=0.19$; HVLT-R total score: $Z=-0.03$, $p=0.98$, $r=0.007$; The Brief Visuospatial Memory Test – Revised (BVMT-R) delayed recall: $Z=-0.27$, $p=0.79$, $r=0.06$; BVMT-R total score: $t(16)=-1.55$, $p=0.14$, $d=0.38$; Wechsler Adult Intelligence Scale – fourth edition (WAIS-IV) Digit Span total score: $t(16)=-0.43$, $p=0.67$, $d=0.11$; D-KEFS Verbal Fluency 1 (phonemic fluency): $t(16)=-1.66$, $p=0.12$, $d=0.40$; D-KEFS Verbal Fluency 2 (semantic fluency): $t(16)=1.38$, $p=0.19$, $d=0.33$; D-KEFS Verbal Fluency 3 (semantic fluency switching categories): $t(16)=0.20$, $p=0.84$, $d=0.05$].

SAINT protocol development:

**Precise targeting to L-DLPFC-sgACC:** Both the left dorsolateral prefrontal cortex (L-DLPFC) and the subgenual cingulate cortex (sgACC) have been shown to be dysfunctional in major depressive disorder (MDD); L-DLPFC has been reliably found to be hypoactive (1–3), and the sgACC is consistently found to be hyperactive in MDD patients (4–6). In addition to the distinct dysfunction in both the L-DLPFC and the sgACC, resting-state functional magnetic resonance imaging (fMRI) data suggest that patients with MDD exhibit reduced anticorrelation between these two regions, which is thought to reflect reduced indirect inhibitory control of the sgACC from the L-DLPFC (7). Repetitive transcranial magnetic stimulation (rTMS) delivered to the L-DLPFC has been shown to normalize hypoactivity in L-DLPFC (8, 9), reduce sgACC hyperactivity (10, 11) and strengthen indirect inhibitory connections between the L-DLPFC and sgACC (7, 12). Neuroimaging data suggest better clinical outcomes are related to higher magnitude anticorrelations (more negative functional connectivity) between the stimulated region of the L-DLPFC and the sgACC (7). This finding has since been confirmed prospectively in other cohorts (13, 14). We restricted our targeting algorithm to find targets within Brodmann Area (BA) 46, as this area has been shown to have the highest mean anticorrelation to the sgACC (7). A recent TMS-fMRI study used resting-state fMRI to target the region of the L-DLPFC that showed the most negative functional connectivity with the sgACC and showed that stimulation of this region propagated to the sgACC in all participants (15). In comparison, in another study that targeted L-DLPFC using anatomical MRI coordinates (the border of BA9 and BA46), stimulation only propagated to the sgACC in 44% of participants (16). These data suggest that using resting-state fMRI to identify and stimulate the region of the L-DLPFC that is most anti-correlated with the sgACC on an individual basis could increase the efficacy of TMS protocols.

**Accelerated delivery:** Multiple spaced stimulation sessions appear to produce non-linear additive effects. Investigations of rat hippocampal brain slices demonstrate that multiple intermittent theta-burst stimulation (iTBS) trains produce a cumulative effect on dendritic spine enlargement (17–19). Studies in humans that have applied TBS protocols to the motor cortex have shown that 2 spaced stimulation sessions produce longer lasting changes in cortical excitability than a single stimulation session (20, 21). Additionally, prior reports indicate that when continuous TBS (cTBS) is delivered to the left posterior parietal cortex in individuals with hemi-spatial neglect, improvements in visual perception resulting from 1 session last 30-40 minutes, 2 spaced applications last 3-8 hours, 4 sessions last 32 hours, and 8 sessions last 3 weeks (22, 23).

**1800 pulses per session**: 1800 pulses per session was chosen as it is the only pulse-dose that has been explored in a blinded iTBS trial (24). Additionally, 1800 pulses per session has been shown to produce long-lasting changes in cortical excitability (25) and optimally produce intended cellular changes (26).

**Overall Dose**: 18,000 iTBS pulses were applied each day of the SAINT protocol to match the number of pulses of a 6-week FDA-approved iTBS protocol (27). In total, across the 5 consecutive days of stimulation, 90,000 iTBS pulses were used to match the total number of 10Hz pulses in a 6-week standard rTMS course (28, 29). Given that 90,000 iTBS pulses equates to 5X the number of pulses given in the FDA-approved 6-week iTBS protocol, this approach allowed for assessment of a wider-range dose-response curve.

**Inter-session interval:** Stimulation sessions were delivered hourly (50-minute inter-session interval) based upon evidence from rat hippocampal slices showing that stimulation with iTBS trains at intervals of 50-90 minutes produces a cumulative effect on dendritic spine enlargement, a process involved in synaptic strengthening. In comparison, iTBS trains delivered with inter-session intervals of 40 minutes or less do not have a cumulative effect on dendritic spines (17–

19). Two iTBS sessions delivered via rTMS to the human prefrontal cortex (30) or motor cortex (31) with a 15-minute inter-session interval have been shown not to increase cortical excitability further than a single iTBS session. A recent study also found that L-DLPFC activity was correlated with sgACC activation 10 minutes after rTMS, but the desired anticorrelation between L-DLPFC and sgACC was observed 27 and 45 minutes after rTMS (32), indicating that intersession interval duration may play an important role, and an interval of 50 minutes may be in the optimal range.

**Stimulation intensity:** Stimulation was delivered at 90% resting motor threshold, as it has been demonstrated that theta-burst stimulation applied <100% rMT produces the optimal change in prefrontal cortical excitability (33, 34). A depth correction (35) was applied to the resting motor threshold to adjust for difference in the cortical depth of the individual's functional target compared to the primary motor cortex in order to consistently achieve 90% rMT in the intended target.

<u>Detailed information about the neuropsychological test battery:</u>

**Verbal learning and memory**

The Hopkins Verbal Learning Test – Revised (HVLT-R) was given to assess learning and recall of verbal information. The HVLT-R (36) is a list-learning task with three learning trials, a 20-minute delayed recall, and a recognition paradigm following the delayed recall. There are 6 alternate forms that allow for serial evaluation.

**Visuospatial learning and memory**

The Brief Visuospatial Memory Test – Revised (BVMT-R) was administered to measure learning and memory of visuospatial stimuli. The BVMT-R (37) is a task that requires the participant to learn an array of simple geometric figures over 3 learning trials. There is a delayed recall after 25 minutes and a recognition paradigm following the delay. There is also a copy task following the memory recall and a recognition task. There are 6 alternate forms that allow for serial evaluation.

**Executive Functioning**

Subtests of the Delis Kaplan Executive Function System (D-KEFS) (38) were used to assess executive functioning. The tests used from the D-KEFS were the Trail Making Test, Color-Word Interference, Verbal Fluency, and the Tower Test. The Trail Making Test (5 conditions) was used to measure combined visuomotor and executive functioning including sequencing and cognitive switching. The test also provides measures of visual scanning and motor speed. The Color-Word Interference test (4 conditions) provides a measure of cognitive inhibition and cognitive switching. There are also word reading and color naming trials that provide a measure of reading and color naming speed. The Verbal Fluency test (3 conditions) provides measures of phonemic and semantic fluency, as well as a trial that involves cognitive switching combined with semantic fluency. The Tower Test provides a measure of planning, learning, and problem solving. This test involves building towers matching pictured models, according to a set of rules that must be followed. The test taker must try to build the tower as quickly as feasible with the fewest number of moves possible.

**Attention and Working Memory**

The Wechsler Adult Intelligence Scale-Fourth Edition (WAIS-IV) (39) Digit Span subtest provides a measure of simple attention and working memory. The test requires the test taker to recall strings of numbers presented verbally and then manipulate them into backward order and sequenced orders.

Resting-state fMRI

Participants were instructed to keep their eyes open and their attention focused on a central fixation point, which consisted of a black screen with a white fixation cross. Participants were also instructed to let their minds wander freely and to avoid any repetitive thoughts such as counting.

Scans were collected with a 3x simultaneous multi-slice (i.e. multiband) acquisition echo planar imaging (EPI) sequence, repetition time of 2000 milliseconds, echo time 30 milliseconds, flip angle of 77 degrees, field of view of 230x230, 128×128 voxel matrix, 1.8×1.8 millimeter$^2$ in-plane resolution, 87 horizontal slices, yielding >1.4 million voxels every 2 seconds. A structural anatomical three-dimensional T1-weighted scan was also collected with a 0.9 millimeter$^3$ voxel volume with 256x256x176 voxel dimensions. Head motion of participants was effectively minimized with the use of memory foam and inflatable padding. Participant alertness during the resting state task was monitored using in-scanner video cameras.

The resting-state scans were spatially smoothed with a 3 millimeter Gaussian kernel, detrended using a linear model of the global signal (40), and band-pass filtered to preserve the typical resting-state frequencies (0.1Hz-0.01Hz).

Targeting algorithms

The first algorithm used the Spearman correlation coefficient between voxel time series as the linkage measurement. Functional sub-units were defined as all voxel pairs being correlated with each other with a Spearman's correlation coefficient of rho greater than or equal to 0.5. Each participant's L-DLPFC was subdivided into a number of functional subunits. This same process was repeated for the bilateral sgACC, with the hierarchical agglomerative clustering algorithm defining the size and shape of each functional subunit based on the correlation coefficient patterns within the bilateral sgACC. For each functional subunit in the L-DLPFC and each subunit in the bilateral sgACC, a single time-series value was created by finding the single voxel time series that was most correlated with the median time series of the subunit. Once a single time series was identified for each subunit, all of the Spearman correlation coefficients were calculated between the L-DLPFC functional subunits and bilateral sgACC subunits to form a correlation matrix. A second algorithm was then deployed to choose the optimal L-DLPFC subunit. The decision-making algorithm considers the net correlation/anticorrelation amount for each L-DLPFC subunit with the bilateral sgACC. This value is calculated using the sum of all the correlation coefficients multiplied by all the sizes of the bilateral sgACC subunits. The decision-making algorithm also considers the size of the L-DLPFC subunit (larger clusters are easier to target with the TMS coil) and the spatial concentration of voxels that make up the subunit. The spatial concentration value was generated by calculating the average of all the three-dimensional Euclidean distances between each of the voxels that make up a subunit and dividing the subunit voxels size by the Euclidean distance measure. The net anticorrelation, the L-DLPFC subunit size, and the spatial concentration of the subunits were the 3 parameters contributing to the decision-making algorithm.

### Supplementary Tables

Supplementary Table 1: Participants' current medications at the time of SAINT

| Participant | Current Medications |
|---|---|
| 1 | Fluoxetine (Prozac) 20mg, amphetamine salts 20mg, quetiapine (Seroquel) 25mg, zolpidem (Ambien) 6.25mg |
| 2 | None. |
| 3 | Dextroamphetamine-amphetamine (Adderall XR) 30mg, alprazolam 0.25-0.5mg/day, vortioxetine (Trintellix) 20mg, zolpidem 10mg, propranolol 30mg |
| 4 | Fluoxetine (Prozac) 40mg |
| 5 | Duloxetine 120mg, amlodipine 5mg, liothyronine 25mcg, cyanocobalamin 1000mcg, Folic acid 400 mcg, zolpidem 10 mg (every other night) |
| 6 | Prozac 20mg and bupropion (Wellbutrin) 300mg |
| 7 | Lamotrigine 200mg, Levothyroxine 100mcg, Liothyronine 10mcg |
| 8 | Bupropion (Wellbutrin) 522mg, dextroamphetamine-amphetamine (Adderall), lisinopril |
| 9 | Cymbalta 120mg and alprazolam (Xanax) 2mg |
| 10 | Lamotrigine (Lamictal) 175mg |
| 11 | Bupropion (Wellbutrin XR) 450mg |
| 12 | None. |
| 13 | Venlafaxine XR 225mg, risperidone 0.5mg, [pantoprazole 20mg][1] |
| 14 | None. |
| 15 | Bupropion (Wellbutrin SR) 400mg, Cytomel 5mcg, Hydroxyzine (Atarax) 50mg, [Flexeril 10mg, doxylamine succinate, estradiol 0.0375mg, Tirosint 50mcg, moxifloxacin HCL, ondansetron 8mg, oxycodone HCL 10-650mg, propranolol HCL 10mg] |
| 16 | Fluoxetine 20mg, atorvastatin 20mg, diazepam 5mg-15mg |
| 17 | Seroquel 300mg, Lexapro 20mg |
| 18 | Bupropion (Wellbutrin XR) 450mg, dextroamphetamine-amphetamine (Adderall) 10mg, vortioxetine (Trintellix) 20mg, ramelteon (Rozerem) 8mg, zolpidem (Ambien) 6.25mg. [atorvastatin (Lipitor) 10mg, hydrochlorothiazide (Hydrodiuril) 0.5mg, lisinopril (Prinivil) 10mg, pantoprazole (Protonix) 40mg, acetaminophen (Tylenol) 500-1000mg, cetirizine (Zyrtec) 10mg, fluticasone propionate (Flonase) 100 mcg, melatonin 5mg] |
| 19 | Vortioxetine (Trintellix) 20mg, alprazolam (Xanax) 0.25mg, [Simvastatin 10mg, estradiol 0.1mg] |
| 20 | Aripiprazole (Abilify) 10mg |
| 21 | None. |

[1]Brackets indicate non-psychiatric medications.

Supplementary Table 2: Clinical assessment scores 1 month after SAINT; mean (SD, n) or %(n)

| | **Pre aiTBS** | **Month post aiTBS** | **Responders (%)[1]** | **Remission (%)[2]** |
|---|---|---|---|---|
| **MADRS** | 34.86 (5.29, 21) | 10.95 (11.76, 20) | 70.00 (20) | 60.00 (20) |
| **HDRS-17** | 25.90 (4.79, 21) | 8.05 (8.31, 20) | 75.00 (20) | 65.00 (20) |
| **HDRS-6** | 13.90 (2.45, 21) | 4.40 (4.72, 20) | 75.00 (20) | 70.00 (20) |
| **BDI** | 28.78 (11.68, 18) | 12.25 (13.06, 16) | 57.14 (14) | 62.50 (16) |
| **C-SSRS[3]** | 1.42 (0.96, 19) | 0.0 (0.00, 19) | 100.00 (14) | 100.00 (19) |
| **HDRS-Q3** | 1.38 (0.67, 21) | 0.10 (0.31, 20) | 100.00 (18) | 90.00 (20) |
| **MADRS-Q10** | 2.38 (0.80, 21) | 0.35 (0.75, 20) | 90.00 (20) | 80.00 (20) |

[1]Response defined as ≥50% reduction in score

[2]Remission defined ≤7 on HDRS-17 (41), ≤4 on HDRS-6 (42), ≤10 on MADRS (43), ≤12 on BDI(44), C-SSRS=0 (45), HDRS-Q3=0 and MADRS-Q10=0

[3]Suicidal ideation subscale

Supplementary Table 3: Clinical assessment scores at 1 month, excluding participants with a treatment history of standard TMS non-response; mean (SD, n) or % (n)

| | **Pre aiTBS** | **Month post aiTBS** | **Responders (%)[1]** | **Remission (%)[2]** |
|---|---|---|---|---|
| **MADRS** | 34.47 (5.85, 15) | 7.07 (7.71, 14) | 85.71 (14) | 71.43 (14) |
| **HDRS-17** | 25.27 (4.91, 15) | 5.57 (6.39, 14) | 85.71 (14) | 78.57 (14) |
| **HDRS-6** | 13.87 (2.33, 15) | 2.64 (3.13, 14) | 92.86 (14) | 85.71 (14) |
| **BDI** | 28.75 (11.78, 12) | 8.27 (10.52, 11) | 77.78 (9) | 72.73 (11) |
| **C-SSRS[3]** | 1.38 (.65, 13) | 0.00 (0.00, 13) | 100.00 (10) | 100.00 (13) |
| **HDRS-Q3** | 1.60 (0.51, 15) | 0.07 (0.27, 14) | 100.00 (14) | 92.86 (14) |
| **MADRS-Q10** | 2.60 (0.74, 15) | 0.36 (0.74, 14) | 92.86 (14) | 78.57 (14) |

[1]Response defined as ≥50% reduction in score

[2]Remission defined ≤7 on HDRS-17 (41), ≤4 on HDRS-6 (42), ≤10 on MADRS (43), ≤12 on BDI(44), C-SSRS=0 (45), HDRS-Q3=0 and MADRS-Q10=0

[3]Suicidal ideation subscale

Supplementary Table 4: Remission rates immediately following the end of treatment for other interventions for TRD

| **Intervention** | **Open Label** | **Blinded** | |
|---|---|---|---|
| | | Active | Placebo |
| Ketamine | 31% (46) | 32% (47) | 4% (47) |
| DBS* | 30% (48) | 18% (49) | 7% (49) |
| ECT | 48% (50) | N/A | N/A |
| MST | 30-40% (51) | N/A | N/A |
| rTMS | 37.1% (52) | 16% (53) | 5.7% (53) |

*DBS outcomes improved as time-from-initiation increased and are reported after 1-year of initiation of treatment.

DBS: Deep brain stimulation, ECT: Electroconvulsive therapy, MST: Magnetic seizure therapy, rTMS: Repetitive transcranial magnetic stimulation.

11. Loo CK et al. High (15 Hz) and low (1 Hz) frequency transcranial magnetic stimulation have

different acute effects on regional cerebral blood flow in depressed patients. *Psychol. Med.* 2003;33:997–1006.

21. Goldsworthy MR, Pitcher JB, Ridding MC. The application of spaced theta burst protocols induces long-lasting neuroplastic changes in the human motor cortex. *Eur. J. Neurosci.* 2012;35(1):125–134.
22. Nyffeler T, Cazzoli D, Hess CW, Müri RM. One session of repeated parietal theta burst stimulation trains induces long-lasting improvement of visual neglect. *Stroke* 2009;40(8):2791–2796.
23. Cazzoli D et al. Theta burst stimulation reduces disability during the activities of daily living in spatial neglect. *Brain* 2012;135:3426–3439.
24. Li CT et al. Efficacy of prefrontal theta-burst stimulation in refractory depression: A randomized sham-controlled study. *Brain* 2014;137(7):2088–2098.
25. Peinemann A et al. Long-lasting increase in corticospinal excitability after 1800 pulses of subthreshold 5 Hz repetitive TMS to the primary motor cortex. *Clin. Neurophysiol.* 2004;115(7):1519–1526.
26. Volz LJ, Benali A, Mix A, Neubacher U, Funke K. Dose-dependence of changes in cortical protein expression induced with repeated transcranial magnetic theta-burst stimulation in the rat [Internet]. *Brain Stimul.* 2013;6(4):598–606.
27. Blumberger DM et al. Effectiveness of theta burst versus high-frequency repetitive transcranial magnetic stimulation in patients with depression (THREE-D): a randomised non-inferiority trial. *Lancet* 2018;391(10131):1683–1692.
28. Janicak PG, Professor of Psy-chiatry is, Carpenter L. The Efficacy of Transcranial Magnetic Stimulation for Major Depression: A Review of the Evidence. *Psychiatr Ann* 2014;44(6):284–292.
29. Modirrousta M, Meek BP, Wikstrom SL. Efficacy of twice-daily vs once-daily sessions of repetitive transcranial magnetic stimulation in the treatment of major depressive disorder: A retrospective study. *Neuropsychiatr. Dis. Treat.* 2018;14:309–316.
30. Chung SW, Rogasch NC, Hoy KE, Fitzgerald PB. The effect of single and repeated

prefrontal intermittent theta burst stimulation on cortical reactivity and working memory [Internet]. *Brain Stimul.* 2018;11(3):566–574.

40. Macey PM, Macey KE, Kumar R, Harper RM. A method for removal of global effects from

fMRI time series. *Neuroimage* 2004;22(1):360–366.

50. Heijnen WT, Birkenhäger TK, Wierdsma AI, Van Den Broek WW. Antidepressant pharmacotherapy failure and response to subsequent electroconvulsive therapy: A meta-

analysis. *J. Clin. Psychopharmacol.* 2010;30(5):616–619.

51. Cretaz E, Brunoni AR, Lafer B. Magnetic Seizure Therapy for Unipolar and Bipolar Depression: A Systematic Review. *Neural Plast.* [published online ahead of print: 2015]; doi:10.1155/2015/521398

52. Carpenter LL et al. Transcranial magnetic stimulation (TMS) for major depression: A multisite, naturalistic, observational study of acute treatment outcomes in clinical practice [Internet]. *Depress. Anxiety* 2012;29(7):587–596.

53. Sehatzadeh S et al. Unilateral and bilateral repetitive transcranial magnetic stimulation for treatment-resistant depression: a meta-analysis of randomized controlled trials over 2 decades. *J. Psychiatry Neurosci.* 2019;44(3):151–163.
